# Predictive Motor Control Based on a Generative Adversarial Network

**DOI:** 10.1101/2023.01.17.524156

**Authors:** Movitz Lenninger, Whan-Hyuk Choi, Hansol Choi

## Abstract

Predictive processing models suggest that a brain decides actions through inference using its internal generative model over the world’s states and their transitions. Most of the predictive processing models have been formalized using explicit representations of the probability distributions, with explicit structures and parameters. They are difficult to learn in general and needs explanation about representation of structure and parameters of distributions, and the method for statistical arithmetic on them, each of which are questions not easy to answer. In this study, we explore an alternative representation for predictive processing which is based on an implicit model known as generative adversarial networks, which has been widely explored recently in machine learning studies as they can learn a distribution directly from data. We demonstrate how a generative adversarial network can be trained to learn an implicit generative model of motor dynamics. And then, we show that such a model can perform approximate inference using the trained model, providing the necessary computations for both the forward and inverse model of motor control. Our framework may provide another formalization for brain’s inference model, especially for learning process. Additionally, we suggest that the functional architecture of the cortical-basal ganglia circuit may modeled as the generator and discriminator in the generative adversarial network model.

## 1 INTRODUCTION

In Neuroscience, there exists a widely influencing idea that brains are inference machines. They use internal generative models to explain the cause of the observations and decide actions to maximize adaptability. The models under the idea are collectively known as predictive processing (Clark, 2012). In the core of the models are the generative model (Pickering and Clark, 2014; Botvinick and Toussaint, 2012; Solway and Botvinick, 2012; Friston et al., 2009), which can be understood as “stochastic recipes for generating observed data” (Gershman, 2019). For example, a probabilistic distribution 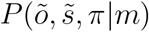 is a generative model to give the probability of observing a sequence of observation *õ*, when the sequence of hidden states and the policy are 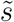 and *π*. The distribution function is parameterized with *m*. The perception is formalized to infer 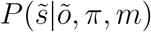, which is the posterior probability of 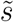 when the probability of a sequence of observation *P* (*õ*) and the agent’s policy *P* (*π*) is given. The action is to find *P* (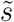, *π*|*õ, m*).

The predictive processing needs to explain how to use the generative models for inference mechanisms. Although the exact posterior can be computed using Bayes rule *P* (*s*|*o*) = *P* (*o*|*s*)*P* (*s*)*/* Σ_*s*_ *P* (*o*|*s*)*P* (*s*), it is intractable for many of realistic conditions when the state *s* is high dimensional, or its functional form is not easy to integrate. The approximate inference methods alleviate the problems. For example, the active inference framework uses a variational Bayes method (Friston, 2010; Smith et al., 2022). It tries to find an approximate distribution *Q*(*s*) of *P* (*s*|*o*), of which the KL-divergence *KL*(*Q* ∥ *P*) = − 𝔼_*Q*_ log(*Q/P*) is minimized. The method can keep the inference procedure tractable by carefully choosing the function *Q*. Active inference demonstrated the methods to approximate 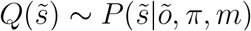 for state perception on the observations and policy, and 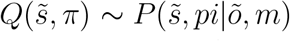 for action generation on observation. The method also provided an update rule for *m* to explain the (parametric) learning of the 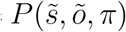 to update the generative model. Studies have demonstrated that predictive processing can explain a wide variety of brain functions, including perception problems such as visual perception or binocular rivalry, or decision-making (Hohwy et al., 2008; Friston et al., 2009). One question is how to implement the statistical arithmetic in explicit model from neural network.

The previous studies use explicit generative models that the probability of samples are explicitly represented in probabilistic distribution functions. (Gershman and Beck, 2017; Gershman, 2019; Bogacz, 2017; Friston et al., 2009). The (approximate) inference is to compute the parameters of the distribution functions using Bayesian or statistical techniques. One question is the representation of explicit generative models in the brains and its inference mechanisms. Although several hypotheses, such as population coding, explain the representation of the probabilities in brains, it is not an easy-to-answer question (Pouget et al., 2013). Moreover, the result of inference is a probability of states or actions. A state or action of the agent must be finally sampled from the distribution. It means that the agent’s state and the generative models should exist independently from each other.

The last important question is learning from the environment. Most predictive processing studies is that the learning are about inference with the model given a prior or to parametric learning. Structural learning of the generative model, for example, learning the functional family of the distribution, is an important task for the brain but mostly overlooked in predictive processing model’s studies.

Implicit generative models are worth investigating as an alternative formalization for predictive processing. The implicit models have a function *G*(*z*^*′*^), which transforms a random sample *z*^*′*^ from an arbitrary distribution *z*^*′*^∼ *P*_*z*_(*z*) to vector *x* in a sample space. The density of the sample *x* = *G*(*z*^*′*^) implicitly represents the *x*’s distribution density. A recently developed machine learning technique called generative adversarial network (GAN) has been successful in training implicit generative models for diverse domains of high dimensional data.(Goodfellow et al., 2014; Radford et al., 2015) Moreover, GAN can learn representation of the data to encode the semantic information. For example,when GAN is trained for face images latent states were found to to encode face directions.(Yeh et al., 2016)

Here, we will demonstrate predictive processing with an implicit generative model based on GAN. First, using a simple toy environment, we show how GAN can learn a generative model of motor system dynamics. Second, we demonstrate how approximate inference using the trained GAN can behave as two of the most important models for motor control: the forward and inverse models. The forward model predicts the system’s future state from the current state and action, and the inverse model decides the actions required to achieve a target state (Pickering and Clark, 2014; Jordan and Rumelhart, 1992; Desmurget and Grafton, 2000; Franklin and Wolpert, 2011). Finally, we will discuss the neural implications of the GAN models. We argue that the functional architecture of the Cortico-Basal Ganglial system can be interpreted in terms of the generator-discriminator system of our GAN predictive processing model.

## 2 MATERIALS AND METHODS

### 2.1 Implicit Generative Model and predictive processing

#### 2.1.1 Planning as inference

Planning is to select a policy *π* which maximizes an objective function such as return. An agent with planning as inference uses an internal generative model 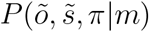 for planning. (Smith et al., 2022). Active inference agents use a prior preference distribution *p*(*o* | *C*), which denotes the agent’s preference for observing a certain observation *o*. The agent infers current hidden state and then infer action with under the current observation and the preferred observation as evidences. The action is biased toward achieving states by the preferred observation distribution.

One of the simplest examples is a single-step state-action-state condition. A state *s* and action *u* is followed by a *s*^*′*^ in a discrete-time system. The generative model for a predictive processing agent is *P* (*o, s, u, o*^*′*^, *s*^*′*^). When we set a pooled hidden state *z* = (*s, s*^*′*^) the generative model is *P* (*o, u, o*^*′*^, *z*). Planning is to infer the *P* (*u*|*o, o*^*′*^), when the current observation is, and the preferred observation is ^*′*^. Bayes rule and marginalization gives *P* (*u* | *o, o*^*′*^) Σ_*z*_ *P* (*o, o*^*′*^| *z*)*P* (*u* | *z*)*P* (*z*) as the exact inference result (Under POMDP setup, *o, o*^*′*^, *u* are independent each other given *z*.). An agent can approximately decide latent states by finding optimal 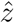 to maximize *P* (*o, o*^*′*^|*z*)*P*(*z*), then, use *P* (*u*|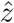) to decide action.

Another important problem in motor control called the forward problem can be also formalized as inference. It is to infer *P* (*o*^*′*^|*o, u*), the future state from the current state in action. Similarly, it is to decide 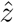 maximizing *P* (*o, u*|*z*)*P* (*z*) and *P* (*o*^*′*^|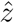) to infer the future observation.

#### 2.1.2 Learning an implicit generative model by GAN

The agents in the real world cannot directly access the generative process of the world, which determines the states of the sample space. Instead, they should learn a generative model from the observations *x* ∼ *p*_*data*_, where *p*_*data*_ is the real distribution density from the generative process. GAN is developed to learn directly from the data. Two functions are learned simultaneously. One is a generator, *G*(*z*; *θ*_*z*_), which maps a random noise *z* with prior distribution *p*_*z*_(*z*) to the sample space and parameterized by *θ*_*z*_ (Goodfellow et al., 2014; Arjovsky et al., 2017; Yeh et al., 2016). Another is *D*(*x*; *θ*_*D*_) called discriminator, which maps a value in sample space to a scalar value. During the training, the purpose of *G* is to generate realistic samples. while *D* plays an adversarial role in discriminating between the motor sequence generated from *G* and the real data. *G* and *D* are trained by optimizing the loss function:

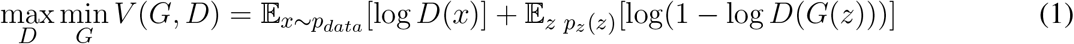

#### 2.1.3 Planning as inference using an implicit generative model

Inference using an implicit generative model is not easy and has not been studied much. For example, Monte Carlo sampling can be used on the implicit generative model. Assume we want to decide an action *u* when current and target observation *oando*^*′*^ are given. We can sample *G*(*z*) using *z* ∼ *P*_*z*_(*z*) and filter them by *O* = *o, O*^*′*^ = *o*^*′*^. The *U* from the filtered samples may provide the target distribution. The method will require to generate a lot of samples, memory to store them, and a mechanism to filter samples and count the samples for density. Moreover, we will need a decision process for action from the filtered samples. We do not want agent’s states independently exist its distribution.

We will investigate a method from machine learning studies. Yeh et al. ((Yeh et al., 2016)) demonstrated how to infer using the GAN generative model. Their problem was to in-paint an omitted part *x*_*omitted*_ of an image when the other part, *x*_*evidence*_ is given. The problem is to find *x*_*omitted*_ maximizing *P* (*x*_*ommited*_|*x*_*evidence*_). When *z* is the latent state of the generative model *P* (*x, z*), the problem is to find *z* which maximizes *P* (*x*_*evidence*_|*z*)*P* (*z*) similar to the previous section. (*x*_*evicence*_ is independent from *x*_*ommited*_ given *z*).

Yeh et al. used GAN, the implicit generative model. Their main idea was to find an optimal input 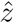 which minimizes a combination of prior loss, *L*_*p*_, and context loss, *L*_*c*_ by gradient-descent.

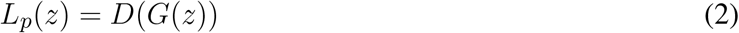

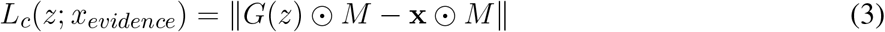

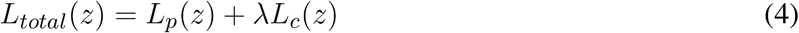

*G* is the generator and *D* is the discriminator. They were pre-trained for the same domain data with the in-paint problems. *x* is a sample with an omitted part. *M* is mask vector for the *x*_*evidence*_ that, *M*_*i*_ = 1 if *x*_*i*_ is given as evidence, *M*_*i*_ = 0, otherwise.

The context loss evaluates the difference between a generated sample *G*(*z*) and the given part of a sample *x*_*evidence*_. As the likelihood *P* (*x*_*evidence*_|*z*) in an explicit form evaluates how well a latent state *z* explains a piece of evidence *x*_*evidence*_, *L*_*p*_(*z*) evaluates related property to the likelihood. (Further studies are needed for the exact relationship). The discriminator evaluates *z* whether a sample *G*(*z*) exists in the trained data in another loss function *L*_*p*_. It measures quantity related to the prior *P* (*G*(*z*))in the explicit Bayesian method, which provides the probability of *G*(*z*) in the overall distribution. Together, the Bayesian posterior maximization for *z* where *P* (*z* | *x*_*evidence*_) ∼ *P* (*x*_*evidence*_ | *z*)*P* (*z*) is related to Yeh’s method, even though further studies are needed for finding the exact relationship between them.

### 2.2 Simulation

#### 2.2.1 A Toy Environment

We will use an arm environment to demonstrate our idea in the following. We will use a simple two-joint model with a single time step (Fig1a), which 14 dimensions. And also, we additionally tested four joint and two-two joint arms models to scale up the model. (See supplementary material) Each has 24 and 28 dimensions.

**Figure 1.**
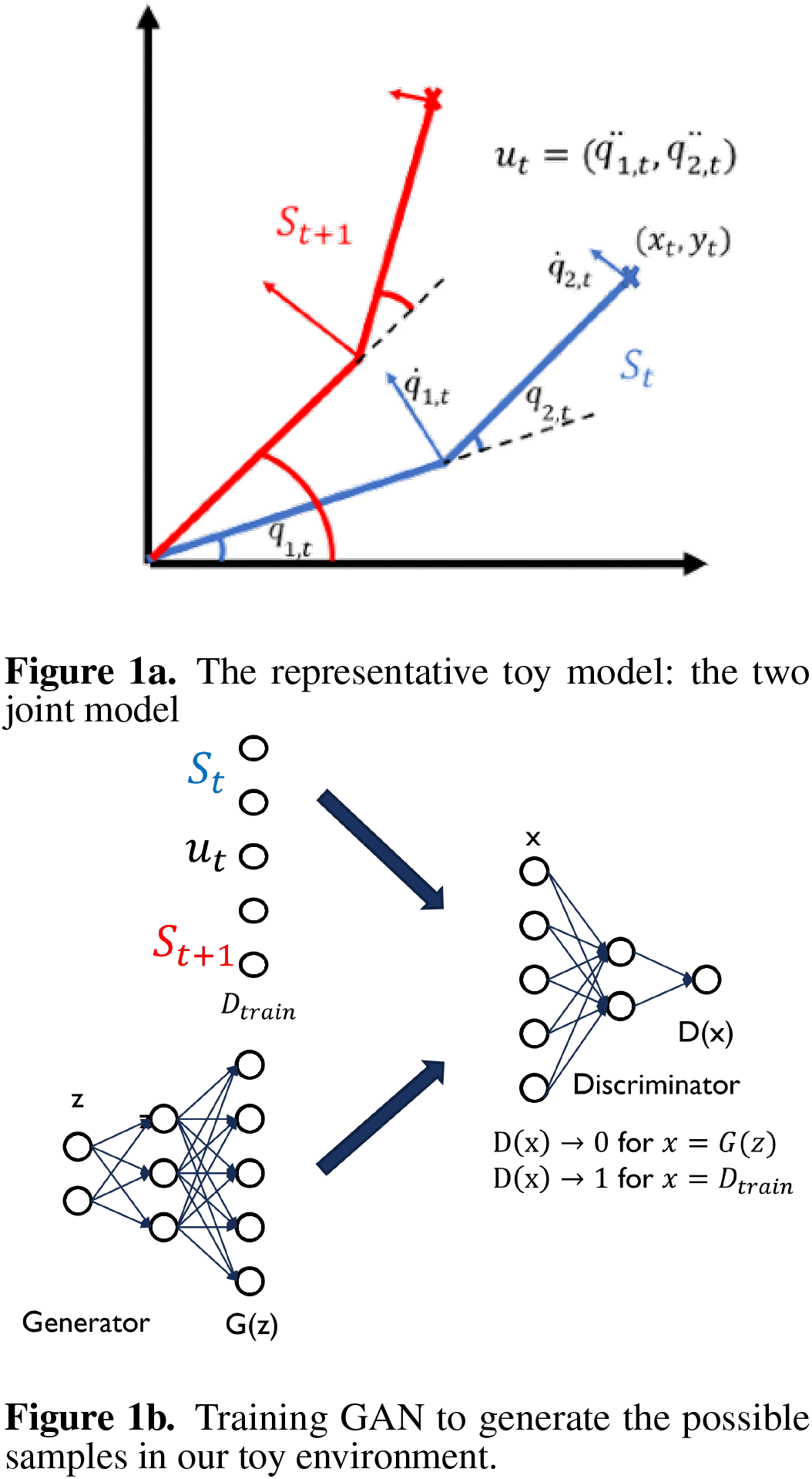
Overall architecture of our study. Our GAN model learns the possible samples in our toy environment. The learnt information is used for action by inference

The variables of our toy arm environment are as below.

- *Q* = (*q*_1_, *q*_2_) *∈* ℝ^2^: The angle of the first and second joint.
- 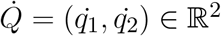: the angular velocities of the joints.
- *U* = (*u*_1_, *u*_2_) *∈* ℝ^2^: *u*_*i*_ the action given on the *i*-th joint.
- (*x, y*) *∈* ℝ^2^: The position of the cursor at the end of the arm.
- 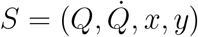: The state of the arm.
- symbols with ^*′*^: The state after action.

The dynamics of the system are as follows:

- 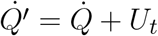
- 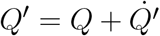

In total, the toy environment has six degrees of freedom. We will use the notation 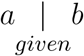 to indicate a, which is fixed from b. For example,

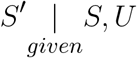

represents the future state, *S*^*′*^ of our toy environment when a set of current states and actions, *S* and *U*, are given.

#### 2.2.2 Training GAN and inference

We uniformly sampled current joint states and the current actions. The joint angles were between [0, *π/*2], angular velocities between [− *π/*6, *π/*6] and the actions between [− *π/*6, *π/*6]. Each dimension was independently normalized between -1 and 1 by first subtracting each dimension’s average and then dividing by the maximum deviation from the average. Two fully connected multilayer perceptrons with leaky ReLU activation functions were used as *G* (6-8-14 nodes) and *D*(14-8-1 nodes; Fig. 1b and trained for the samples from toy environment. The forward problem is to infer *S*^*′*^ when the current state *S* and action *U* are given. The inverse problem is to infer an action, *U*, which brings the cursor to a desired position (*x, y*) from the current state *S* is given. The approximate inference method using GAN explained above was used to infer the target values.

## 3 RESULTS

### 3.1 Training Generative Model for a Two-Arm System

We trained a GAN generative model on data generated from the toy environment. Fig. 2a shows representative samples from the naive and trained model, respectively. During the training process, we evaluated the GAN by letting it generate 1000 random inputs **z** and tested whether the *S*^*′*^ in *G*(*z*) is close to 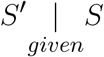, *U* where *S* and *U* from *G*. Comparing the joint states and cursor positions between the actual and generated future states shows that as the number of iterations increases, the model-generated samples get closer to obeying the dynamics of the actual toy model environment. Both the error between the actual and generated future joint states and the error between the actual and generated cursor position decrease during training (Fig. 2b). This demonstrates that the adversarial training process is capable of producing a generative model which reasonably reflects the actual dynamics of the environment solely based on training data.

**Figure 2.**
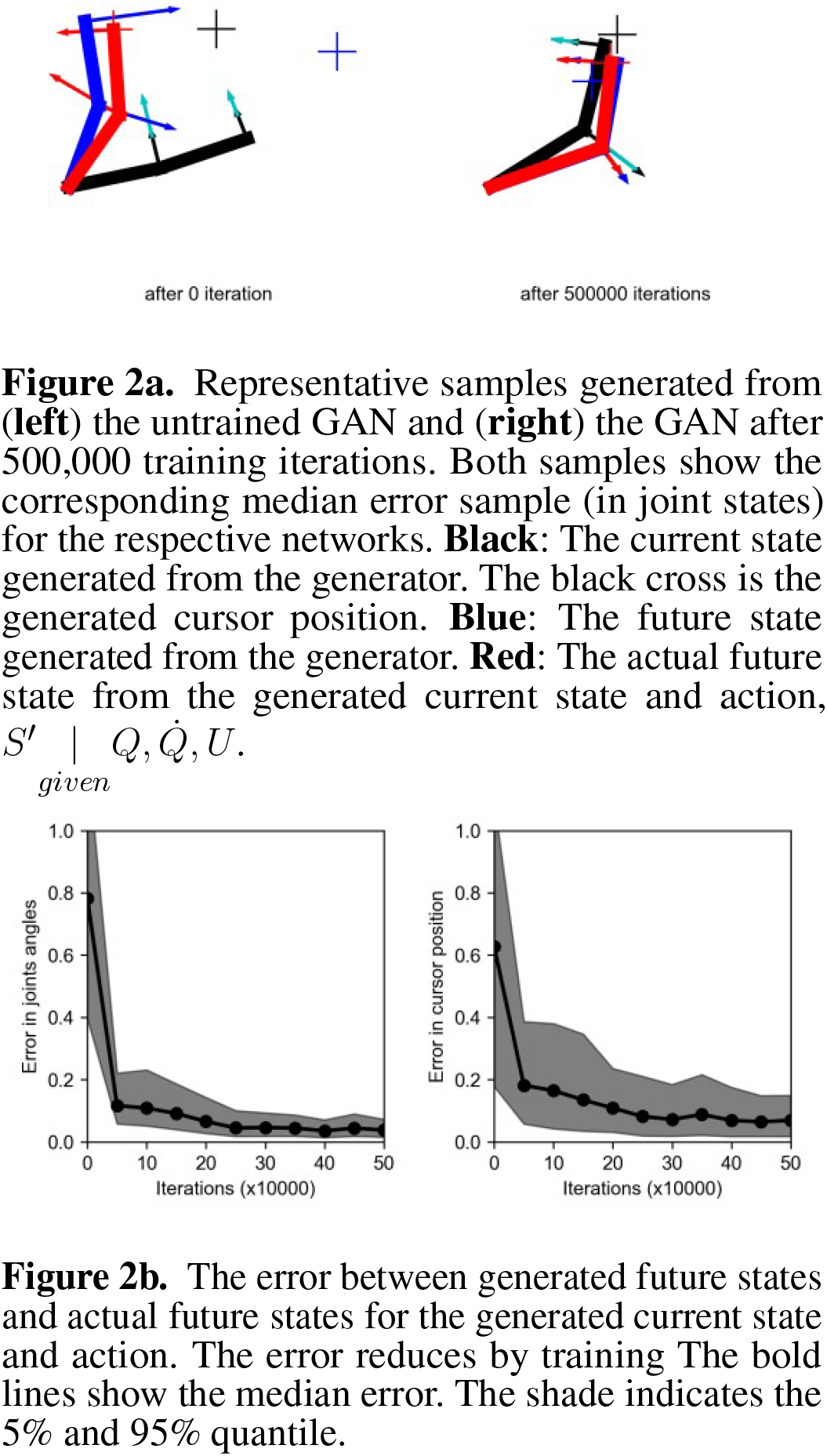
Training a Generative Model. GAN could internalize the dynamics of our toy model that, it could generate the samples, which are close to realistic samples after training

### 3.2 The Forward and Inverse Model by Approximate Inference with GAN

To investigate whether the trained GAN model can be used as a forward model, we generated 100 test samples from the toy environment and masked the future states. Based on the knowledge of the current state and action(Fig. 3, black arms), the generative model is tasked with completing the sample for the prior and context loss. Compared to the naive model, the trained model is much better at predicting the future state of the arm (compare blue and red arms in Fig. 3a). The error for the future joint states between the actual and predicted future state (joint angle and angular velocities) decreases with increasing iterations (Fig. 3b). The same holds for the error between the predicted position of the cursors from the forward model and the actual future cursor position (Fig.3b).

**Figure 3.**
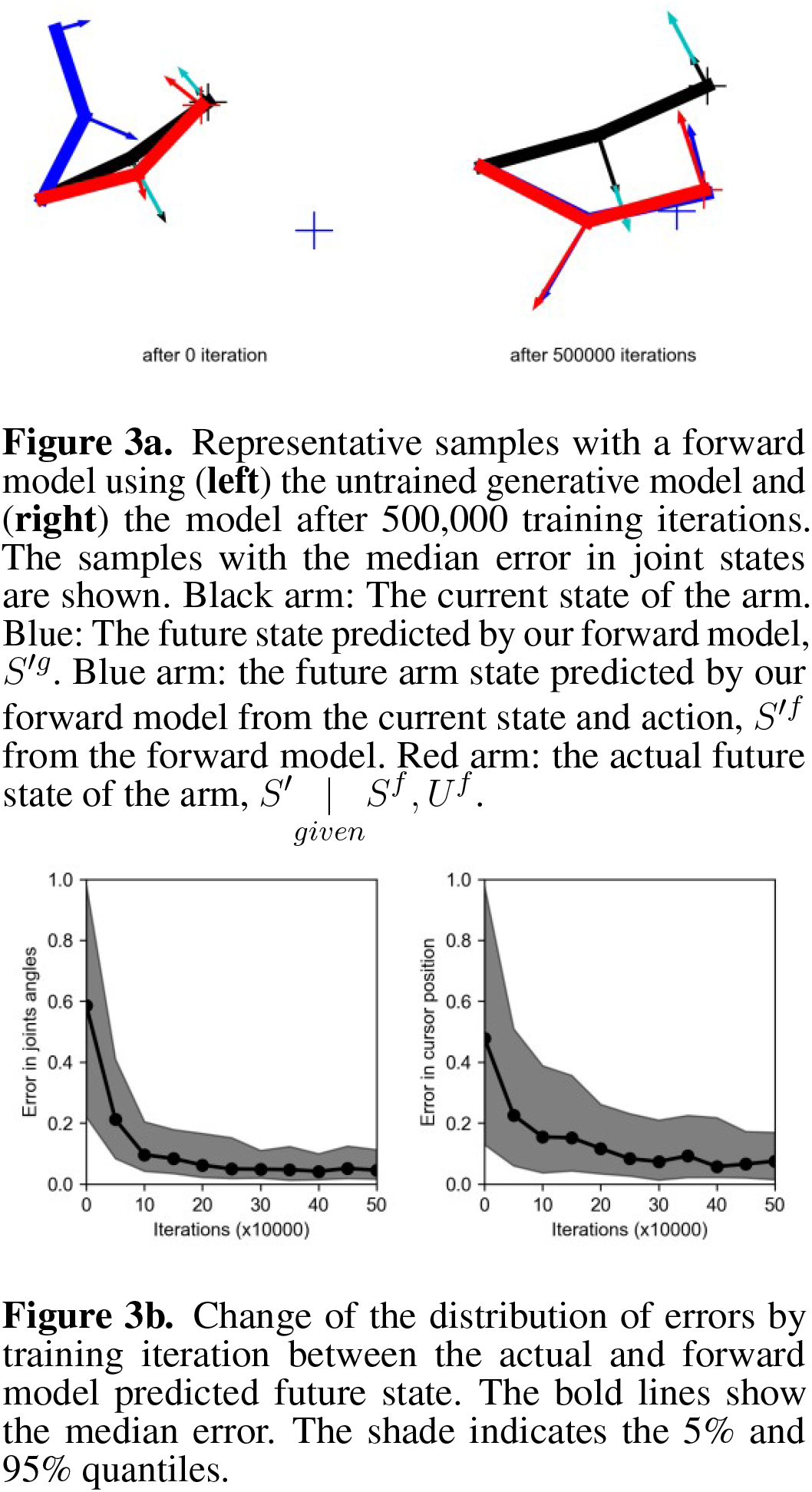
The forward model. Using the trained GAN network for the toy model samples, the future state of an arm could be inferred from the current state and action. The error between inferred and real future states were reduced by the training.

The same approximate inference method is used as an inverse model (Fig. 4). In the inverse model, the task is to predict the action when the agent’s current state and target cursor positions are known (black arms and dots in Fig. 4a, respectively). As the generative model trains, the inferred action by the inverse model becomes more accurate compared to the untrained model, that it moves the cursor closer to the target position (red crosses in Fig. 4a).

**Figure 4.**
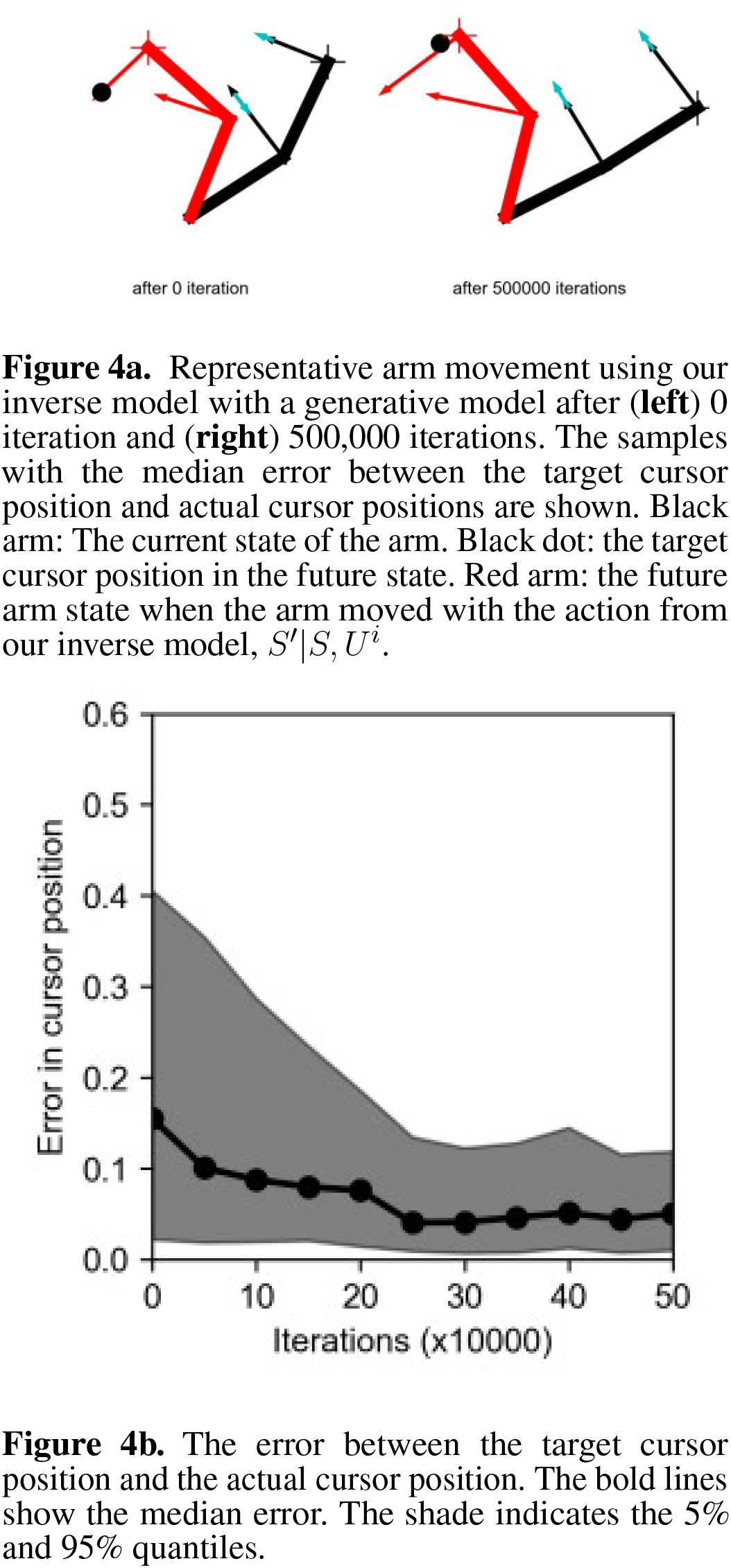
The inverse model. Using the trained GAN network for the toy model samples, the action could be inferred from the current state and target cursor state. The inferred action could move the cursor closer to the target as the iteration increase.

We tested our system on two more conditions. One with a case with non-task related dimensions that two two joints arms exist and only one arm is related with task (supplementary fiigure 2). The other was with four joints arm (supplementary fiigure 2). They had more dimensions to learn the generative models (28 and 24 dimensions correspondingly). For the both of the cases, our system could learn the generative model and be used to infer the better action to achieve target condition compared to the naive system.

## 4 DISCUSSION

### 4.1 Implication in neuroscience

We hypothesize that this functional architecture of GAN for implicit generating model and inference method can be related to the functional architecture of the Cortico-Basal Ganglial (BG) system: the neocortex as the generator and the BG as the discriminator.

Firstly, the discriminator requires (1) to receive signals from different sources of inputs to discriminate actual motion from internally generated, (2) to be capable of reducing the high-dimensional inputs to a lower-dimensional evaluation value, and (3) to send the evaluation signal to both the generator and the discriminator to teach and instruct.

The Basal Ganglia (BG) system may have those desired characteristics. (1) The dense input neurons of BG receive inputs from the diverse areas of the brain (Gurney et al., 2001; Bar-Gad and Bergman, 2001; Bar-Gad et al., 2003; Watabe-Uchida et al., 2012), especially from the neocortex and BG reduces the dimensions of the inputs to the activity of the smaller dimension of DAergic neurons (Bar-Gad et al., 2003; Watabe-Uchida et al., 2012). (2) We suggest that the phasic DAergic signal may act as the discriminator signal. Phasic DAergic activity has been suggested to function as a temporal difference signal (i.e., the difference between the predicted and arrived state value) for reinforcement learning (Montague et al., 2004; Schultz et al., 1997; Houk et al., 1995). Importantly, recent studies suggest alternative roles of DA neurons to represent non-reward information (Tian et al., 2016; Noudoost and Moore, 2011) and state prediction errors (Wei et al., 2015; Engelhard et al., 2019; Schwartenbeck et al., 2016; Suarez et al., 2019; Takahashi et al., 2017; Nakahara et al., 2004). Gardner et al. (Gardner et al., 2018) suggested that the phasic DA signal may not only encode the reward prediction error but perhaps act as a general prediction error signal in a diverse feature space. The prediction error can be interpreted as an evaluation signal of the generated samples, as suggested by Pfau and Vinaylas (Pfau and Vinyals, 2016). (3) The DA projects back to the basal ganglia as well as to diverse brain areas, including the neocortex (see (Puig et al., 2014) for a review). Thus, the discriminatory signal can be accessible to both the discriminator and generator networks. These functional structures of dimensional reduction, encoding of low-dimensional evaluation signals about how realistic a sample is, and signal projections of the evaluation signals are all necessary (even though not sufficient) properties for a discriminator.

The generator requires (1) to receive evaluation input from the discriminator and to be trained from the signal, (2) to generate potential samples for the control sequences, and (3) to map abstract latent input into a specific sample. We propose that the neocortex may have the necessary properties.

(1) The neocortex receives DAergic projections from the BG, which produce synaptic triad (Yao et al., 2008) and modifies the synapse by the prediction error signal. The DAergic prediction error signal may teach the neocortex area. (2) The motor cortex activity is correlated with the current state and actions (Fried et al., 2011; Cisek and Kalaska, 2005; Li et al., 2016; Churchland et al., 2010, 2012) as well as imaginary motor movement (Schnitzler et al., 1997; Doud et al., 2011). The imagination of motor movement is one way of motor sample generation, which is a possible behavior by a generator. (3) The cortical area is suggested to have a functional hierarchy ranging from abstract higher causes to more concrete sensory-motor states for visual perception (DiCarlo et al., 2012), motor control (DeWolf and Eliasmith, 2011; Tani, 2016; Dumoulin et al., 2017), and decision making (Ribas-Fernandes et al., 2011, 2018).

The place for the forward and inverse model inference process is also needed. First of all, the generator needs to generate samples matching the context cues. The affordance competition (Pezzulo and Cisek, 2016; Cisek and Kalaska, 2005; Cisek, 2007) suggests a process of contexts limit the sample space. A signal of target states can activate the neurons correlated with the potential actions to achieve the target (Cisek and Kalaska, 2005). When multiple actions are possible for the target signals, the neurons correlated with competing actions are activated together, and the neural correlates for the irrelevant actions for the given context are reduced.

Secondly, another criterion to generate actions in our model is prior loss, in which the discriminator evaluates the samples generated by the generator by a trained model. We hypothesize that the thalamic output from BG may have the necessary properties. The BG projects to the thalamus as one of the circuit’s outputs. Interestingly, the thalamic output of BG has been suggested to be correlated with decision-making among multiple choices (Dunovan and Verstynen, 2016; Wei et al., 2015; Gurney et al., 2015; Keuken et al., 2015). This evaluation is not only limited to the value of the decision but also related to prior decision-making(Ding and Gold, 2013; Lak et al., 2017). With the function of DA as a general prediction error signal explained above, we suggest that the BG might function as a general evaluator for generated samples based on the reward size and the probability of the sample.

### 4.2 Conclusion

In this study, we suggested a predictive processing model using an implicit generative model. GAN was capable of training a generative model for our toy environment motor control sequences. The training lowered the error between the generated samples and the real constraints of the toy environment. Importantly, although minimal structural assumptions were imposed beforehand, the system could work adequately after training in the environment. The forward and inverse models of motor control were constructed as approximate inference processes as data in-painting processes. Interestingly, the same approximate inference architecture was used for both models. The forward and inverse models only differed in what information is considered the evidence and what information is omitted. We suggested the potential relationship between the explicit Bayesian statistics and the context and prior loss that the context loss is related to the likelihood of *P* (*o* | *z*) latent state for the observation and the prior loss as the prior for *P* (*G*(*z*)) or *P* (*z*). Clearly, our suggestion remains conceptual and further mathematical and experimental studies are required.

We focused on the functional architecture of the system. How can the generated samples’ evaluation signal teach the generator and discriminator? Our GAN framework for modeling brain functions needs a lot more clarifications. For example, it uses back-propagation in the learning and inference process, which is still a topic of debate and may not be feasible in real neural systems. Another algorithm, such as the recently suggested forward-forward algorithm (Hinton, 2022), may provide alternative options but still needs further studies.

Another limitation is from the simple one-step generative model; the generative cannot explain the complex behavior requiring multiple steps of choice. Further studies are needed, such as a hierarchical generative model or successor representation, which represents the multiple steps of data as the density of states.

#### 4.2.1 Permission to Reuse and Copyright

Figures, tables, and images will be published under a Creative Commons CC-BY license, and permission must be obtained for the use of copyrighted material from other sources (including re-published/adapted/modified/partial figures and images from the internet). It is the responsibility of the authors to acquire the licenses, follow any citation instructions requested by third-party rights holders, and cover any supplementary charges.

## CONFLICT OF INTEREST STATEMENT

The authors declare that the research was conducted in the absence of any commercial or financial relationships that could be construed as a potential conflict of interest.

## AUTHOR CONTRIBUTIONS

ML designed, implemented code, analyzed data, and wrote the manuscript. WC analyzed the result and wrote the manuscript. HC designed, analyzed data, wrote the manuscript, and supervised the project.

## FUNDING

Whan-Hyuk Choi is supported by the National Research Foundation of Korea(NRF) grant funded by the Korean government (NRF-2022R1C1C2011689).

## ACKNOWLEDGMENTS

We thank Prof. Dr. Carsten Mehring, University of Freiburg, for the comments that greatly improved the manuscript.

## Notes

### Competing Interest Statement

The authors have declared no competing interest.

### Summary of Updates

Added supplementary materials with a few more simulation results

