## supplementary_material for "Predictive Motor Control Based on a Generative Adversarial Network"

frontiers

### Supplementary Material

#### 1 SUPPLEMENTARY DATA

We tested two additional toy models to test how our GAN based motor control system behave in more complex conditions. One is to scale up the model to four joints. The environment has four joints arm. It's sample is now 24 dimensions (10 dimensions for current and future states each: four for angle, four for angular velocity and two for the cursor position. four dimensions for action). Another is to test whether our model can recognize the responsible dimensions when the toy models containing non-responsible dimensins for target actions. To test this, we built a toy model with two two-joint arms. The models were trained for the both of the arms but the action inference was only related with one of them.

#### 2 SUPPLEMENTARY TABLES AND FIGURES

##### 2.1 Figures

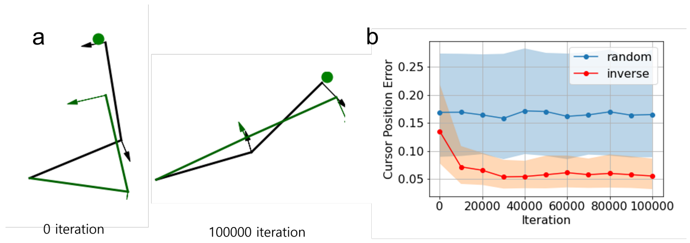


**Figure S1.** Two two-joint arms condition shows the selection of dimension, responsible for the task. **(A)** representative movement of an arm by our model inferred action, after 0 (left) and 100000 (right) iterations of training. Black arm is the current and the green dot is the target state. Green is the future state after action from our system. Only the right arm, which is the responsible for this task is shown. **(B)** The difference between the target position and the position of cursor after moving arm. As the training iteration increase, it decreases.

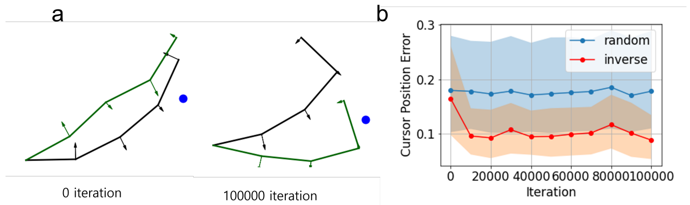


**Figure S2.** Four joint arm system shows that our system works for scaled-up model. **(A)** representative movement of an arm by our model inferred action, after 0 (left) and 100000 (right) iterations of training. Black arm is the current and the green dot is the target state. Green is the future state after action from our system. **(B)** The difference between the target position and the position of cursor after moving arm. As the training iteration increase, it decreases.
